# A nucleus-vacuole junction in fission yeast enriches the HMG-CoA reductase Hmg1 and INSIG protein Ins1

**DOI:** 10.64898/2026.05.18.725815

**Authors:** Akika Murayama, Shintaro Fujimoto, Yasushi Tamura

## Abstract

Membrane contact sites (MCSs) enable communication between organelles and play central roles in lipid metabolism. In budding yeast, the nucleus-vacuole junction (NVJ) functions as a dynamic platform that integrates lipid metabolism and stress responses. However, it remains unclear whether NVJ structure and function are broadly conserved across eukaryotes, particularly because Nvj1, the key membrane tethering factor that mediates NVJ formation in budding yeast, is absent in higher eukaryotes. Here, we investigated whether an MCS analogous to the NVJ in budding yeast exists in fission yeast (*Schizosaccharomyces pombe*), which lacks Nvj1. We show that an NVJ is present in fission yeast and serves as a platform for the accumulation of sterol synthesis factors, including the HMG-CoA reductase Hmg1 and the INSIG homolog Ins1. We further demonstrate that the localization of these factors depends on the membrane protein insertase Snd302 and is dynamically regulated by nutrient conditions. Our findings reveal that, despite the absence of Nvj1, the NVJ is functionally conserved as a site for sterol synthesis in fission yeast, suggesting a conserved role of spatial organization in lipid metabolism.

## Introduction

Membrane contact sites (MCSs) are regions where membranes of distinct cellular compartments come into close proximity, facilitating inter-organelle communication and serving as important sites for lipid metabolism (Calí et al., 2025; Prinz et al., 2020; Reinisch et al., 2025; Tamura et al., 2019; Voeltz et al., 2024). Among these, the nucleus-vacuole junction (NVJ) in budding yeast (*Saccharomyces cerevisiae*) is known as a platform that coordinates lipid metabolism and cellular responses to nutrient conditions (Henne and Hariri, 2018; Kohler and Büttner, 2021; Kvam and Goldfarb, 2006; Malia and Ungermann, 2016). In budding yeast, the NVJ is formed primarily through the interaction between the nuclear envelope protein Nvj1 and the vacuolar membrane protein Vac8 (Jeong et al., 2017; Pan et al., 2000), whereas Mdm1 also contributes to NVJ formation independently of Nvj1/Vac8 (Hariri et al., 2019; Henne et al., 2015). In addition to these core tethers, multiple lipid transport proteins localize to the NVJ, including Osh1, an oxysterol-binding protein homolog (Levine and Munro, 2001); Nvj2, an SMP domain-containing protein (Liu et al., 2017); Ltc1 (Lam6), a VASt/StART-like domain-containing protein (Murley et al., 2015); and Vps13, a bridge-like lipid transfer protein (Lang et al., 2015). These proteins harbor lipid transport-related domains, suggesting that the NVJ serves as a platform for lipid exchange between organelles. In addition, lipid biosynthetic enzymes including the enoyl reductase Tsc13, are also concentrated at this site (Kohlwein et al., 2001). Another NVJ-associated factor, Nvj3, has been linked to Dga1-dependent triacylglycerol synthesis and lipid droplet biogenesis at ER contact regions (Adebayo et al., 2026). The recruitment of Tsc13, Osh1, and Nvj2 to the NVJ depends on Nvj1, whereas Nvj3 is targeted to this site in an Mdm1-dependent manner.

Glucose starvation (GS) induces NVJ expansion and remodeling, accompanied by the accumulation of sterol synthetic enzymes and regulatory factors, including the HMG-CoA reductases Hmg1 and Hmg2 and the INSIG homologs Nsg1 and Nsg2 (Fujimoto and Tamura, 2026; Hugenroth et al., 2025; Rogers et al., 2021). Concomitantly, GS is associated with reduced activity of fatty acid elongases such as Elo2 and Elo3, which may promote the recruitment of the aspartyl protease Ypf1 to the NVJ, where it is required for the recruitment of sterol regulatory factors (Fujimoto and Tamura, 2026). In addition, Snd3, a membrane protein insertase of the SRP-independent (SND) pathway, is recruited to the NVJ upon GS and promotes NVJ expansion downstream of PKA and Snf1 signaling (Aviram et al., 2016; Tosal-Castano et al., 2021; Yang et al., 2025). In contrast, the peroxisome-associated protein Pex31 is enriched at the NVJ under glucose-replete conditions and restricts GS-induced NVJ remodeling, as its loss leads to constitutive NVJ expansion and recruitment of starvation-specific factors even in the presence of glucose (Hugenroth et al., 2025). Furthermore, Msc1, which localizes to the perinuclear space between the inner and outer nuclear membranes and is strongly induced under GS, mediates this GS-dependent NVJ remodeling (Mito et al., 2026). The NVJ also serves as the site of piecemeal microautophagy of the nucleus, a selective autophagic process in which portions of the nucleus are directly engulfed by the vacuole (Roberts et al., 2003). These observations highlight the NVJ as a dynamic regulatory hub that integrates lipid metabolism, nutrient signaling, and organelle quality control.

Despite extensive characterization in budding yeast, it remains unclear whether NVJ structure and function are conserved across other eukaryotes. Notably, Nvj1, the central tethering factor required for NVJ formation in budding yeast, is not conserved in fission yeast (*Schizosaccharomyces pombe*) or in higher eukaryotes. This raises a question as to whether an NVJ-like MCS exists in these organisms. In addition, the regulatory mechanisms governing sterol metabolism differ markedly between budding and fission yeast. In particular, fission yeast conserves the SCAP-SREBP pathway, a key regulatory system for sterol biosynthesis found in higher eukaryotes (Bien and Espenshade, 2010; Brown et al., 2018; Hughes et al., 2005), whereas budding yeast lacks this pathway and instead relies on sterol-responsive transcription factors such as Upc2 and Ecm22 (Vik and Rine, 2001). These differences suggest that both the spatial organization and regulatory control of sterol biosynthesis may have diverged during evolution. Given these considerations, it remains unknown whether NVJ exists in fission yeast and whether it serves as a platform for sterol metabolism.

In this study, we demonstrate that the NVJ is present in fission yeast and serves as a platform where sterol regulatory factors, including the HMG-CoA reductase Hmg1 and the INSIG homolog Ins1, are enriched. Furthermore, we show that their localization is dynamically regulated by nutrient conditions and depends on the membrane protein insertase Snd302. Together, these findings highlight a conserved mechanism for the spatial regulation of sterol metabolism and provide a framework for understanding how lipid metabolism is spatially organized across eukaryotes.

## Results and Discussion

### A nucleus-vacuole junction is present in fission yeast

The nucleus-vacuole junction (NVJ) has been extensively characterized in budding yeast (*Saccharomyces cerevisiae*), and numerous NVJ-resident proteins (hereafter referred to as ScNVJ proteins) have been identified. Many of these factors are conserved in fission yeast (*Schizosaccharomyces pombe*) (Table 1). However, Nvj1, a central tethering factor critical for NVJ formation in budding yeast, is not conserved in fission yeast or in higher eukaryotes. We therefore investigated whether an NVJ-like membrane contact site (MCS) is present in fission yeast despite the absence of Nvj1. To address this, we analyzed the intracellular localization of fission yeast homologs of ScNVJ proteins (hereafter referred to as SpNVJ candidate proteins). GFP or mNeonGreen was genomically fused to the C-terminus of each candidate protein. Confocal microscopy showed that Hmg1-GFP and Ins1-GFP were localized to the nuclear envelope and formed punctate structures that were frequently positioned adjacent to vacuoles visualized by autofluorescence (Fig. 1A). The autofluorescent signals were confirmed to represent vacuoles by co-labeling with the vacuolar membrane marker Vac8-GFP and FM4-64 (Fig. S1A). To further characterize the spatial relationship between these puncta and vacuoles, vacuole membranes of cells expressing Hmg1-GFP or Ins1-GFP were stained with FM4-64 and imaged by fluorescence microscopy. Both proteins localized to sites where the nuclear envelope was closely apposed to vacuoles (Fig. 1B, Movie 1), indicating the presence of NVJ in fission yeast. Hereafter, NVJ in fission yeast and budding yeast are referred to as spNVJ and ScNVJ, respectively. Hmg1 is the HMG-CoA reductase, a rate-limiting enzyme in ergosterol biosynthesis, whereas Ins1 is an INSIG homolog that negatively regulates sterol synthesis by inhibiting Hmg1 activity through phosphorylation (Burg et al., 2008). Consistently, simultaneous imaging of Ins1-GFP and Hmg1-mCherry revealed their colocalization at these sites (Fig. 1C). Together, these results strongly suggest that NVJ exists in fission yeast and serves as a platform for the spatial regulation of ergosterol biosynthesis.

**Table 1.**
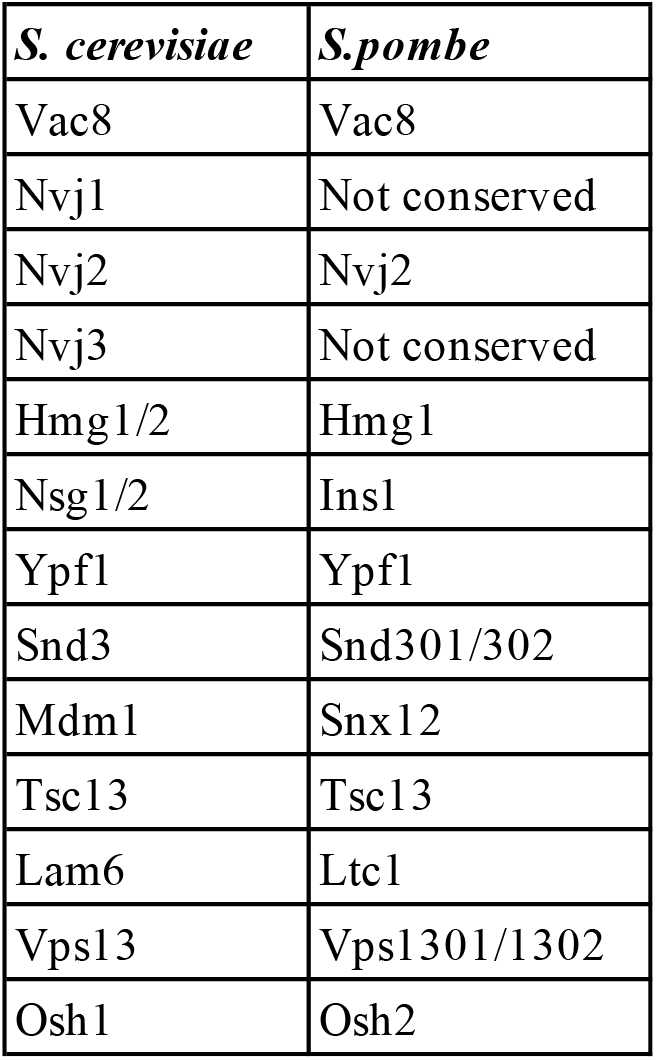
Conservation of budding yeast NVJ-associated factors in fission yeast.

| *S. cerevisiae* | *S.pombe* |
| --- | --- |
| Vac8 | Vac8 |
| Nvj1 | Not conserved |
| Nvj2 | Nvj2 |
| Nvj3 | Not conserved |
| Hmg1/2 | Hmg1 |
| Nsg1/2 | Ins1 |
| Ypf1 | Ypf1 |
| Snd3 | Snd301/302 |
| Mdm1 | Snx12 |
| Tsc13 | Tsc13 |
| Lam6 | Ltc1 |
| Vps13 | Vps1301/1302 |
| Osh1 | Osh2 |

**Fig. 1.**
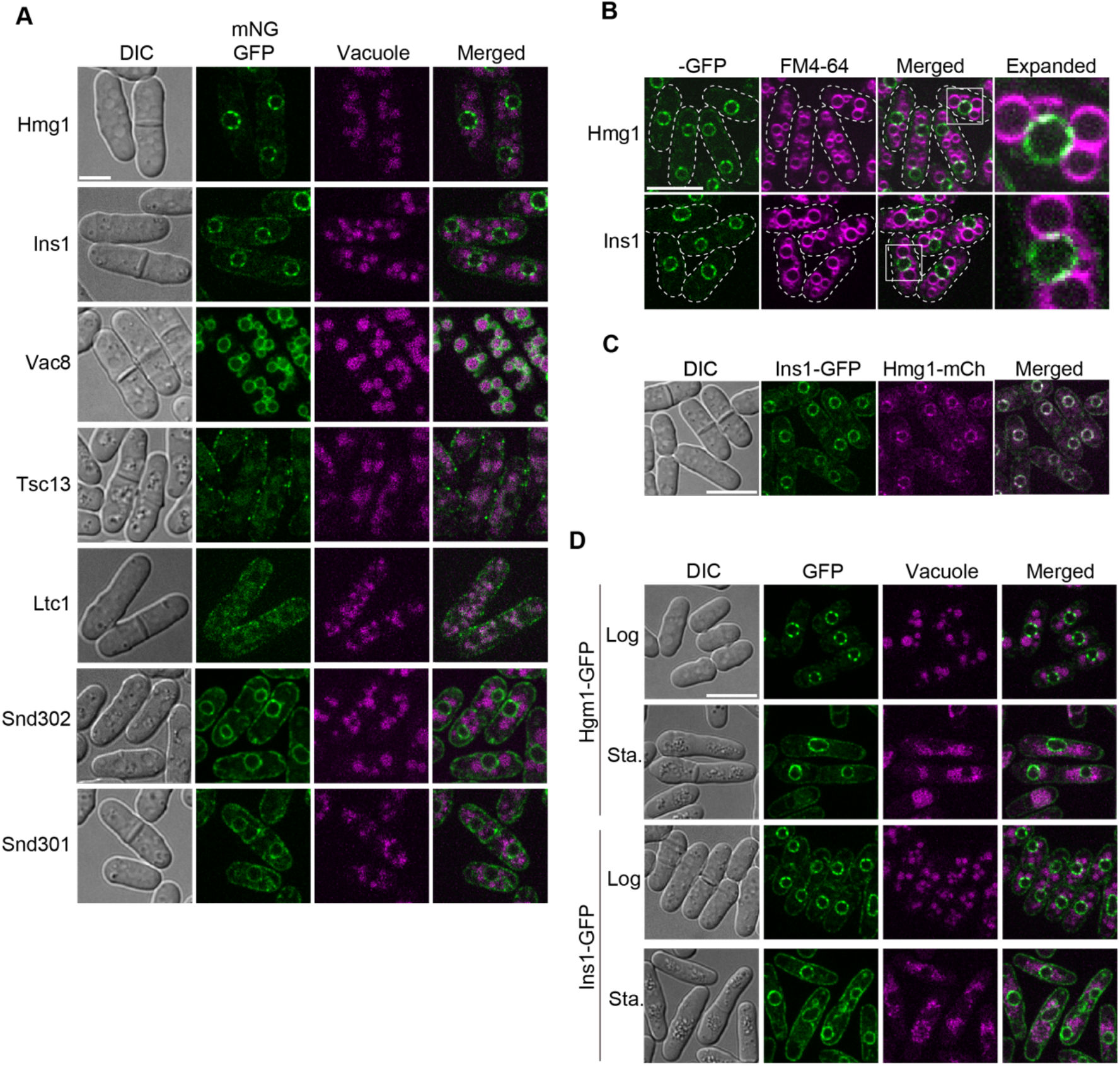
Hmg1 and Ins1 are enriched at the NVJ in fission yeast. (A) Fission yeast orthologs of NVJ-localized proteins in budding yeast were tagged with GFP or mNeonGreen (mNG) and examined by fluorescence microscopy. Scale bar, 5 µm. (B) Cells expressing Hmg1-GFP or Ins1-GFP were stained with FM4-64 to visualize vacuoles and observed by confocal microscopy. Dashed lines indicate the cell boundary. Boxed regions are shown at higher magnification. Scale bar, 10 µm. (C) Confocal images of cells co-expressing Ins1-GFP and Hmg1-mCherry. Scale bar, 10 µm. (D) Cells expressing Hmg1-GFP or Ins1-GFP were observed during logarithmic growth or after 2 days of culture to reach stationary phase. Vacuoles were visualized by autofluorescence. Scale bar, 10 µm.

In fission yeast, Vac8, which interacts with Nvj1 in budding yeast, was distributed throughout the vacuolar membrane and did not exhibit punctate enrichment resembling that of Hmg1 or Ins1 (Fig. 1A). Tsc13 and Ltc1 displayed punctate signals along the cell periphery, suggesting enrichment at ER-plasma MCSs. The Snd3 homologs, Snd301 and Snd302, were broadly distributed along the ER membrane, with occasional puncta observed on the nuclear envelope. Interestingly, approximately 30% of Snd302 puncta were located adjacent to vacuoles, suggesting that Snd302 is partially enriched at the spNVJ. Ypf1, Snx12 (the homolog of budding yeast Mdm1), Nvj2, Osh2, and the Vps13 homologs Vps1301 and Vps1302 did not show clear enrichment at specific membrane compartments (Fig. S1B).

Hmg1, Hmg2 and the INSIG homologs Nsg1 and Nsg2 accumulate at the scNVJ under glucose starvation (GS) conditions (Fujimoto and Tamura, 2026). However, Hmg1 and Ins1 were enriched at the spNVJ during logarithmic growth, whereas in stationary phase they became more broadly distributed across the ER membrane (Fig. 1D). On the nuclear envelope, they formed partial enrichment domains distinct from the punctate spNVJ structures. These observations indicate that spNVJ localization of Hmg1 and Ins1 is regulated by nutrient conditions. The distinct nutrient-dependent patterns of NVJ localization between budding and fission yeast may reflect fundamental differences in metabolic regulation. The budding yeast exhibits strong glucose repression of respiration and preferentially undergoes aerobic fermentation (De Deken, 1966; Pfeiffer and Morley, 2014). During GS, cells undergo a diauxic shift toward respiratory metabolism, coinciding with scNVJ accumulation of Hmg1/2 and Nsg1/2, in budding yeast. In contrast to budding yeast, fission yeast is a petite-negative species that cannot normally survive the loss of mitochondrial DNA (Heslot et al., 1970), indicating a greater physiological dependence on mitochondrial respiration and oxidative phosphorylation. Consistent with this, mitochondrial respiration remains important for rapid cell proliferation in fission yeast even under glucose-rich conditions (Malecki et al., 2016; Malecki et al., 2020). This reliance on respiration during fermentative growth raises the possibility that the NVJ localization of sterol regulatory factors such as Hmg1 and Ins1 is functionally important under conditions that require mitochondrial respiration. However, the mechanistic relationship between respiratory metabolism and NVJ function remains largely unexplored and warrants further investigation.

### Efficient NVJ localization of Hmg1 and Ins1 requires Snd302

We next sought to identify factors required for spNVJ targeting of Hmg1 and Ins1. Loss of Ins1, which colocalizes with Hmg1 at the spNVJ, did not alter the localization pattern of Hmg1, indicating that Hmg1 can target the spNVJ independently of Ins1 (Fig. 2A). In budding yeast, scNVJ recruitment of Hmg1/2 and Nsg1/2 depends on Nvj1 and Ypf1 (Fujimoto and Tamura, 2026). However, loss of the fission yeast homologs of Vac8 or Ypf1 did not impair spNVJ localization of Hmg1 (Fig. 2A). Loss of Vac8 resulted in vacuolar enlargement, which was accompanied by expansion of the spNVJ domain marked by Hmg1 (Fig. 2A). Similarly, deletion of Snx12, the homolog of Mdm1 that contributes to scNVJ formation independently of Nvj1/Vac8 in budding yeast, also caused vacuolar enlargement and apparent expansion of the Hmg1-positive spNVJ domain.

**Fig. 2.**
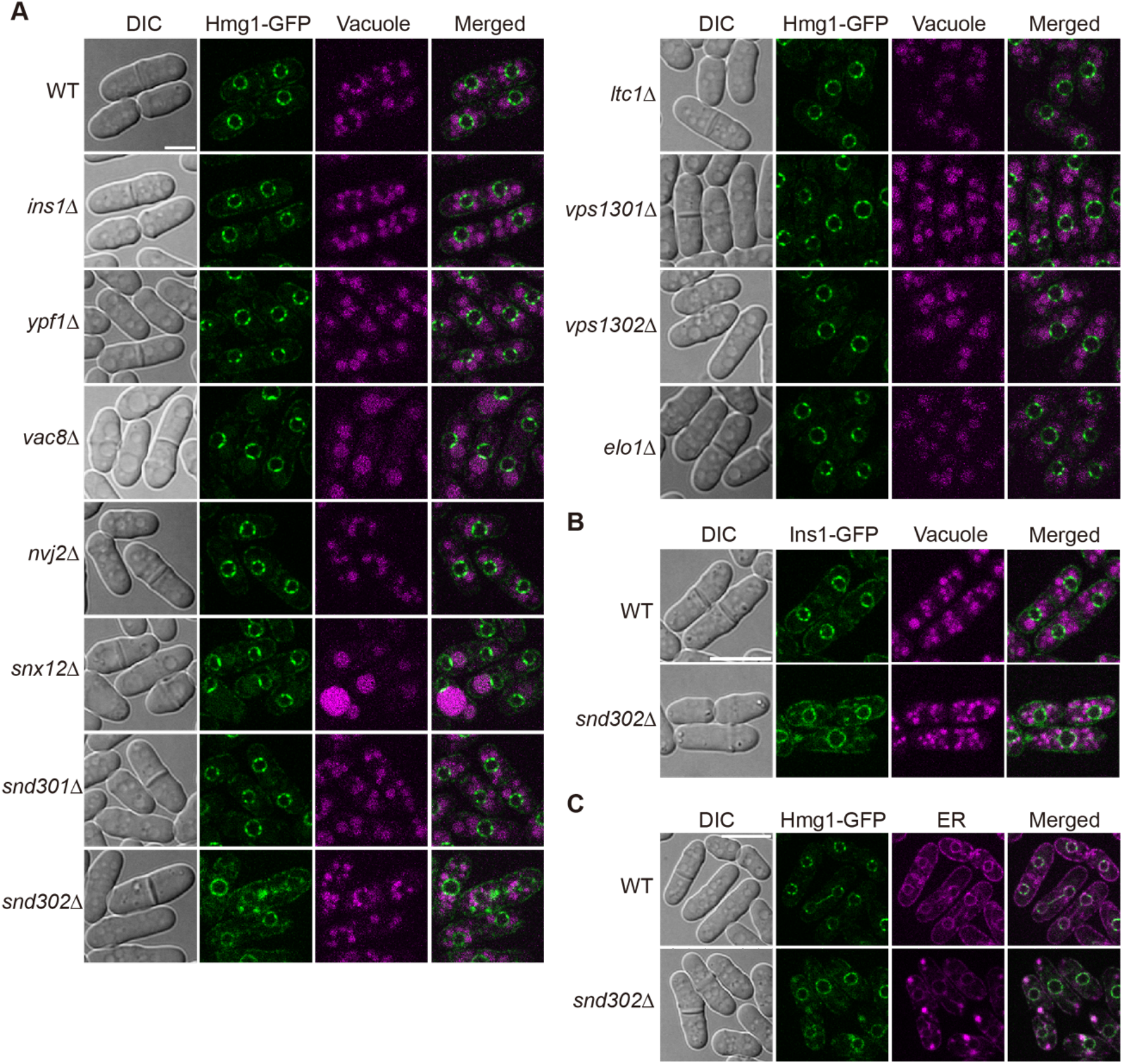
Loss of Snd302 reduces NVJ localization of Hmg1 and Ins1. (A) Confocal images of wild-type and the indicated gene deletion strains expressing Hmg1-GFP. Vacuoles were visualized by autofluorescence. Scale bar, 10 µm. (B) Confocal images of wild-type and snd302Δ cells expressing Ins1-GFP. Vacuoles were visualized by autofluorescence. Scale bar, 5 µm. (C) Localization of Hmg1-GFP and the ER marker BipN-mCherry-ADEL in wild-type and *snd302*Δ cells. Scale bar, 10 µm.

Deletion of Nvj2, Ltc1 (the Lam6 homolog), or Vps1301 and Vps1302 (Vps13 homologs), all of which are associated with lipid transport at the NVJ, likewise had no detectable effect on Hmg1 localization (Fig. 2A). Although we previously found that inhibition of fatty acid elongation promotes GS-specific scNVJ remodeling in budding yeast, we did not observe differences of Hmg1 spNVJ partitioning in cells lacking Elo1, a fatty acid elongase (Fig. 2A). In contrast, loss of Snd302, a homolog of budding yeast Snd3 that accumulates at the scNVJ during GS, abolished spNVJ localization of Hmg1 (Fig. 2A). Instead, Hmg1 became dispersed across the nuclear envelope and ER and additionally formed aberrant spherical structures distinct from vacuoles., A similar phenotype was observed for Ins1-GFP (Fig. 2B). Notably, loss of snd301 another Snd3 homolog, did not show this phenotype, suggesting a specific requirement for Snd302 in spNVJ localization of Hmg1 and Ins1 (Fig. 2A). Because Hmg1 and Ins1 formed spherical structures in *snd302Δ* cells instead of their typical localization patterns along the nuclear envelope and ER, we visualized ER morphology by expressing BipN-mCherry-ADEL, in which mCherry is fused to the N-terminal ER-targeting signal of Bip and the ER retention signal ADEL. These experiments revealed the loss of Snd302 caused pronounced alterations in ER morphology, including the formation of spherical membrane structures (Fig.2C).

Previous studies have shown that the SND proteins constitute an alternative targeting route to the ER, and Snd3 is thought to function as a membrane protein insertase (Aviram et al., 2016; Yang et al., 2025). It is therefore conceivable that Snd3 accumulation at the scNVJ facilitates preferential delivery of specific membrane proteins to this domain under stress conditions. The partial accumulation of Snd302 at spNVJ sites suggests that it may contribute to membrane protein biogenesis within restricted regions of the ER/nuclear envelope. Moreover, disruption of ER morphology in *snd302Δ* cells may result from defective insertion or trafficking of multiple membrane proteins, including those required for ER architecture or lipid biosynthesis.

### Snd301 and Snd302 have redundant functions

Sequence comparison revealed that Snd301 and Snd302 are highly similar, differing only by a modest extension at the C-terminus of Snd302 (Fig. 3A). Despite this similarity, only deletion of Snd302 affected spNVJ localization of Hmg1 and Ins1 (Fig. 2A). To test whether the C-terminal extension confers functional specificity, we generated truncated Snd302 variants by inserting a 3×FLAG tag into the genomic locus after residues 147, 159, or 182, thereby removing the distal C-terminal region. Confocal imaging showed that all truncated mutants showed normal spNVJ localization of Hmg1-GFP, indicating that the C-terminal extension is not required for the spNVJ partitioning of Hmg1 (Fig.3B).

**Fig. 3.**
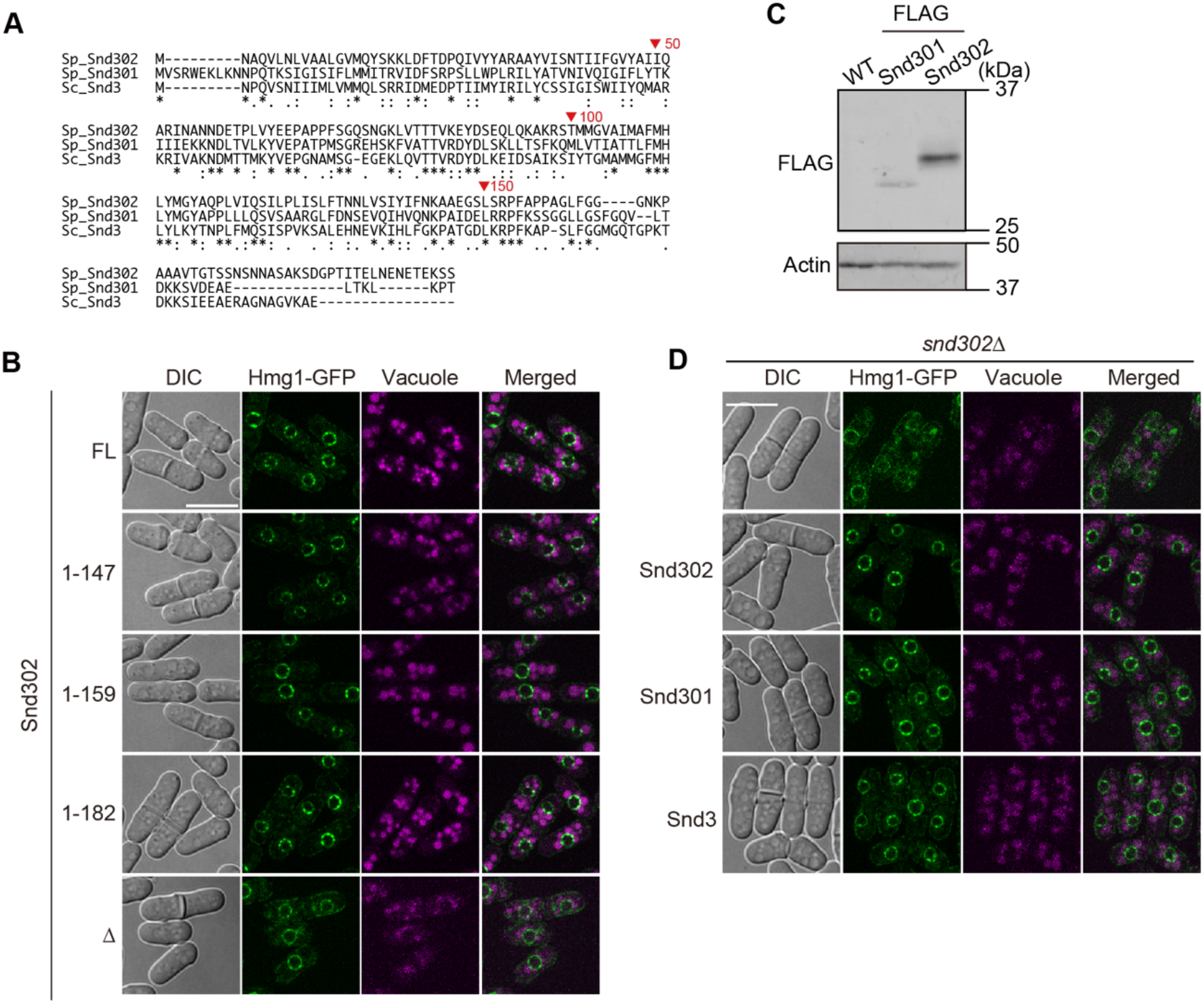
Snd301 and Snd302 have redundant functions. (A) Amino acid sequence alignment of Snd301, Snd302, and budding yeast Snd3. (B) Localization of Hmg1-GFP in cells expressing C-terminally truncated mutants of Snd302. Δ indicates *snd302*Δ cells. Scale bar, 10 µm. Vacuoles were visualized by autofluorescence. (C) Immunoblot analysis of total lysates from cells expressing Snd301-FLAG or Snd302-FLAG. (D) Rescue of Hmg1 localization by overexpression of Snd301, Snd302, or budding yeast Snd3 in *snd302Δ* cells, analyzed by fluorescence microscopy. Vacuoles were visualized by autofluorescence. Scale bar, 10 µm.

We next compared protein abundance by introducing 3×FLAG tags at the endogenous *snd301*^*+*^ and *snd302*^*+*^ loci. Immunoblot analysis revealed that Snd302 is expressed at approximately sixfold higher levels than Snd301 (Fig. 3C). These findings suggest that the apparent functional specificity of Snd302 may result from differential expression levels rather than intrinsic functional divergence between the two proteins. Consistent with this idea, overexpression of Snd301 from the strong *ef1a* promoter (Matsuyama et al., 2008) restored spNVJ localization of Hmg1-GFP in *snd302Δ* cells (Fig.3D). Furthermore, overexpression of budding yeast Snd3 also complemented the snd302 deletion phenotype (Fig.3D). These results strongly suggest that Snd301 and Snd302 share largely redundant functions and that their differential contributions to NVJ organization arise from differences in their expression level.

### Glucose starvation-induced relocalization of Hmg1 depends on Ins1 but not Snd302

scNVJ size and protein composition are dynamically regulated in response to GS (Fujimoto and Tamura, 2026). The spNVJ localization pattern of Hmg1 differed between logarithmic and stationary phases (Fig. 1D), suggesting that spNVJ organization is also responsive to nutrient conditions. We therefore examined whether the localization pattern of Hmg1 at the spNVJ changes in response to GS. Confocal microscopy analysis revealed that, whereas Hmg1 was enriched at the spNVJ during glucose-rich logarithmic growth, approximately 70% of wild-type cells exhibited expanded semicircular Hmg1 structures along the nuclear envelope under GS conditions, and these structures were not always associated with vacuoles (Fig. 4A, C). To further characterize Hmg1 localization, we reconstructed three-dimensional images from confocal z-stacks. During logarithmic growth, Hmg1 exhibited multiple small patch-like structures (Fig. 1A, Movie.1). In contrast, under GS conditions, Hmg1 frequently formed a single large patch along the nuclear envelope (Movie.2). These observations suggest that GS induces not only expansion of the spNVJ-associated Hmg1 domain but also redistribution of Hmg1 to non-spNVJ regions of the nuclear envelope. Interestingly, although loss of Snd302 strongly impaired NVJ partitioning of Hmg1 during logarithmic growth (Fig. 2A), *snd302Δ* cells still exhibited semicircular Hmg1 structures under GS at levels similar to those in wild-type cells (Fig. 4B, C). These findings indicate that GS-induced relocalization of Hmg1 occurs largely independently of Snd302. In contrast, *ins1Δ* cells displayed a decreased frequency of semicircular Hmg1 patterns together with increased diffuse localization along the nuclear envelope, indicating that Ins1 plays an important role in the GS-induced relocalization of Hmg1 (Fig. 4B, C). In budding yeast, GS induces Snf1 (AMP-activated kinase homolog)-dependent NVJ remodeling (Fujimoto and Tamura, 2026). To test whether a similar mechanism operates in fission yeast, we analyzed Hmg1 localization in cells lacking the Snf1 homolog Ssp2 (Matsuzawa et al., 2012). Deletion of Ssp2 impaired GS-induced relocalization of Hmg1 to expanded nuclear envelope domains to a similar extent as observed in *ins1*Δ cells (Fig. 4B, C). These findings suggest that both Snd302 and Ssp2 contribute to GS-dependent relocalization of Hmg1 to expanded nuclear envelope domains.

**Fig. 4.**
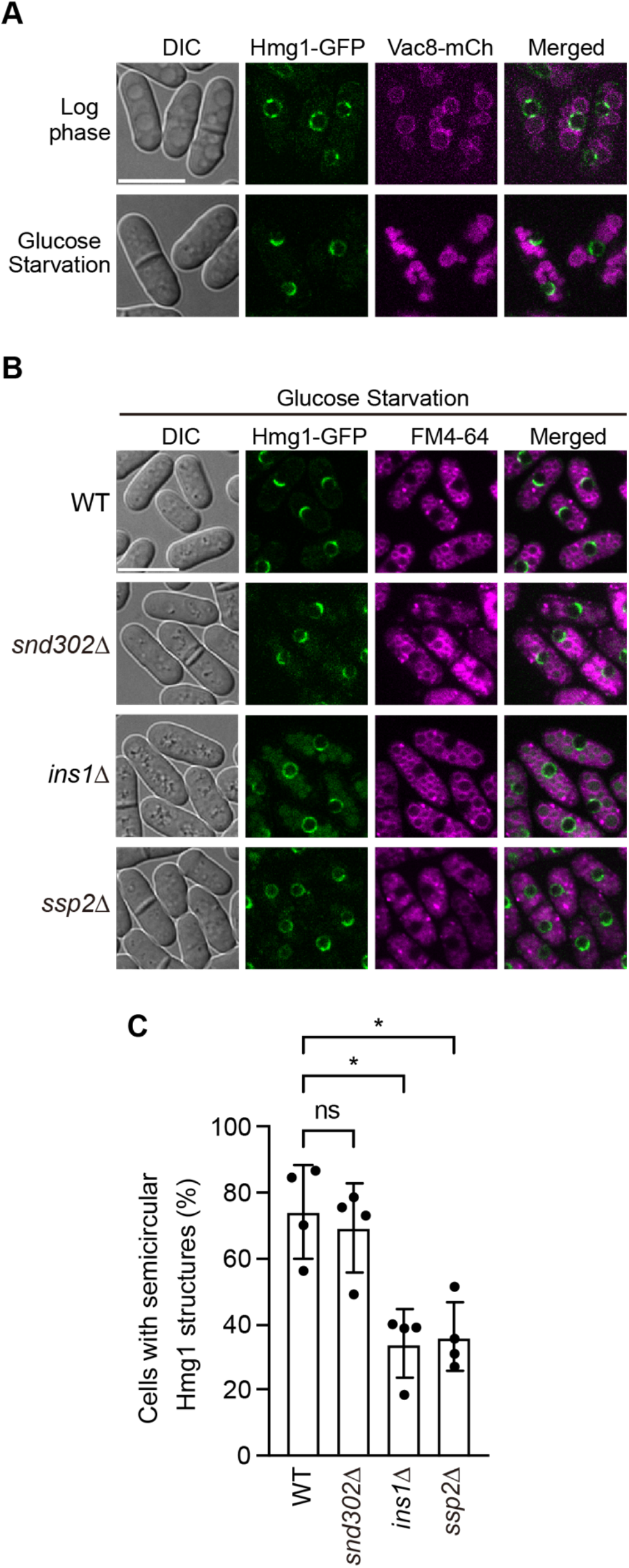
Hmg1 localization dynamically changes in response to GS. (A) Confocal images of cells expressing Hmg1-GFP during logarithmic growth or after glucose starvation. Vacuolar membranes were visualized by co-expression of Vac8-mCherry. Scale bar, 10 µm. (B) Localization of Hmg1-GFP in wild-type, *snd302Δ, ins1Δ*, and *ssp2Δ* cells under glucose starvation conditions. Vacuoles were stained with FM4-64. Scale bar, 10 µm. (C) Bar graph shows the percentage of cells displaying semicircular Hmg1 localization patterns under GS conditions. Values are means ± S.E. from four independent biological replicates. The numbers of cells scored in each replicate were as follows: WT, n = 178, 183, 254, and 272 cells (total = 887); snd302Δ, n = 75, 223, 164, and 114 cells (total = 576); ins1Δ, n = 116, 269, 135, and 265 cells (total = 785); and ssp2Δ, n = 284, 274, 236, and 300 cells (total = 1,094). * indicates P < 0.05 (Mann– Whitney U test).

Taken together, our findings demonstrate that the NVJ is present in fission yeast and serves as a spatial platform for the accumulation of ergosterol regulatory factors, including the HMG-CoA reductase Hmg1 and the INSIG protein Ins1. Efficient targeting of these proteins to the spNVJ requires the membrane protein insertase Snd302, suggesting a link between membrane protein biogenesis pathways and spatial organization of the nuclear envelope. Furthermore, nutrient-dependent relocalization of Hmg1 differs substantially between budding and fission yeast. These findings suggest that although the use of NVJ-related membrane domains for sterol regulation is conserved, the mechanisms governing enzyme localization and spatial organization have diverged between species. This divergence may reflect differences in sterol regulatory systems. Notably, fission yeast retains the SCAP-SREBP pathway conserved in higher eukaryotes (Bien and Espenshade, 2010; Brown et al., 2018; Hughes et al., 2005), whereas budding yeast lacks this system. Thus, analysis of NVJ organization in fission yeast may provide important insights into spatial regulation of sterol metabolism in higher eukaryotes, including humans.

Several important questions remain unresolved. First, the molecular basis of NVJ formation in fission yeast remains unclear. Identification of factors that directly mediate interactions between the nuclear envelope and vacuoles will therefore be an important focus of future studies. In addition, the physiological significance of NVJ-associated sterol metabolism in fission yeast remains poorly understood. In particular, whether Hmg1 targeting to NVJ-related domains directly contributes to sterol biosynthetic output, membrane remodeling, or stress adaptation remains to be determined. It will also be important to clarify whether Snd302 selectively mediates targeting of a specific subset of NVJ proteins or more broadly regulates trafficking of membrane proteins, thereby indirectly influencing localization of Hmg1 and Ins1. Finally, the mechanisms underlying nutrient-dependent redistribution of Hmg1 to non-NVJ nuclear envelope domains during GS remain unclear. Future studies examining how metabolic signaling pathways and respiratory activity influence NVJ organization will provide important insights into how spatial regulation of lipid metabolism is integrated with global cellular physiology.

## Materials and Methods

### Yeast strains and growth conditions

The fission yeast strain AY160-14D (*h90 ade6-216 leu1-32 lys1-131 ura4-D18*), used as the parental strain in this study, was provided by the National Bio-Resource Project (NBRP), Japan (JPNBRP202225) (Hayashi et al., 2009). Fission yeast cells were grown in YES medium (0.5% yeast extract, 3% D(+)-glucose, 225 mg/l adenine, 225 mg/l L-histidine, 225 mg/l L-leucine, 225 mg/l uracil, and 225 mg/l lysine) or in SD-Leu medium (0.67% yeast nitrogen base without amino acids, 2% glucose, supplemented with appropriate amino acids and nucleic acid bases). Gene deletion and tagging were performed by homologous recombination using appropriate gene-targeting cassettes. For gene disruption, DNA fragments containing the clonNAT resistance marker (*natNT2*) were amplified from pBS-natNT2 (Kojima et al., 2019) using primers listed in Table S1 and introduced into fission yeast cells. Transformants carrying the *natNT2* marker were selected on YES plates supplemented with clonNAT at a final concentration of 100 µg/ml. For C-terminal tagging, GFP, mNeonGreen, and 3×FLAG tags were introduced by PCR-based gene targeting using pFA6a-3×FLAG-KanMX, pFA6a-GFP-S65T-KanMX, pFA6a-6xGly-FLAG-hphMX4, or pFA6a-mNeonGreen-3×FLAG-hphMX as templates (Funakoshi and Hochstrasser, 2009; Longtine et al., 1998), together with primers listed in Table S1. Transformants carrying the *kanMX* or hphMX marker were selected on YES plates containing 100 µg/ml G418 or 300 µg/ml hygromycin, respectively. All yeast strains used in this study are listed in Table S2. Glucose starvation medium was prepared by omitting glucose from YES medium.

### Plasmids

Plasmids for overexpression of Snd301, Snd302, and budding yeast Snd3 in fission yeast were constructed as follows. The *snd302*^+^ gene was amplified from fission yeast genomic DNA using primers 6497 and YU6118, and inserted into the NheI/SalI-digested pDUAL-FFH61c vector (Matsuyama et al., 2008) using the SLiCE method (Motohashi, 2015). For Snd301 and budding yeast Snd3, the respective genes were amplified from fission yeast or budding yeast genomic DNA using primer pairs 6771/6772 or 6768/6770, respectively, and cloned into the NheI/SalI-digested pDUAL-Pef1a vector (Matsuyama and Yoshida, 2012) using the SLiCE method (Motohashi, 2015). To generate integration constructs, the resulting plasmids were digested with NotI to remove the *ars1-ura4*^*+*^ fragment. The linearized fragments were introduced into fission yeast cells and integrated into the chromosomal *leu1* locus by homologous recombination.

### Fluorescence Microscopy

Fission yeast cells were grown in 5 ml YES liquid medium at 30°C for 18 h to mid-log phase (OD_600_ ≈ 1.0). Cells were harvested by centrifugation at 2,000 × g for 30 s at 25°C, washed once with 5 ml Milli-Q water, and resuspended in a small volume of Milli-Q water. A 2-µl aliquot was placed on a glass slide (Matsunami, S7441), covered with a coverslip (Matsunami, C218181), and mounted with immersion oil (Olympus, IMMOIL-F30CC). Cells were observed using an IX-83 fluorescence microscope (Olympus) equipped with a CSU-X1 confocal scanner (Yokogawa Electric Corporation) and a 100× objective lens (UPLSAPO100XO, NA 0.75). GFP and mNeonGreen were excited at 488 nm, FM4-64 at 561 nm, and vacuolar autofluorescence at 405 nm using OBIS lasers (Coherent). Z-stack images were acquired at 0.2-µm intervals using an EMCCD camera (Evolve 512, Photometrics). Image processing was performed using ImageJ (NIH). Three-dimensional movies were generated from the z-stack images using NIS-Elements software (Nikon). For vacuolar staining, cells (OD_600_ ≈ 0.2) were incubated with FM4-64 (final concentration 40 µg/ml) at 30°C for 90 min, washed, and further incubated in fresh YES medium for 1 h at 30°C. Cells were then washed with Milli-Q water and prepared for microscopy as described above. GS was induced by collecting cells equivalent to 2 OD_600_ units by centrifugation at 2,000 × g for 30 s at 25°C, resuspending them in 5 ml glucose-free YES medium, and incubating at 30°C for 24 h. Cells were then washed with Milli-Q water and prepared for microscopy as described above.

### Immunoblotting

Total cell lysates of fission yeast were prepared as follows. Cells were grown in 5 ml YES medium at 30°C to OD_600_ ≈ 1.5. Cells were harvested by centrifugation at 2,000 × *g* for 5 min at 25°C, washed once with 800 µl Milli-Q water, and pelleted at 20,000 × *g* for 5 min. The cell pellet was resuspended in 500 µl of 0.3 M NaOH and incubated at 25°C for 5 min. After centrifugation at 20,000 × *g* for 5 min, the supernatant was removed, and the pellet was resuspended in 100 µl of 1× SDS sample buffer and incubated at 95°C for 5 min to generate total lysates. Proteins were separated by SDS-PAGE and transferred onto a PVDF membrane (Millipore, IPFL00010) using a semi-dry blotting system (ATTO, WSE-4045) at a constant voltage of 18 V for 1 h. After transfer, membranes were blocked in 1% skim milk in TBS-T buffer (10 mM Tris-HCl, pH 7.5, 150 mM NaCl, 0.05% Tween-20) for 1 h at 25°C. Membranes were incubated with primary antibodies (anti-FLAG and anti-Actin antibodies)diluted in Can Get Signal Solution I (TOYOBO, NKB-201) overnight at 4°C. After incubation, membranes were rinsed four times with Milli-Q water and washed in TBS-T for 30 min at 25°C. Secondary antibodies (Cy5 AffiniPure Goat Anti-Mouse IgG (H+L) , Jackson Immuno Research Laboratories, Inc. 115-175-146) diluted in Can Get Signal Solution II (TOYOBO, NKB-301) were then applied, and membranes were incubated for 1 h at 25°C in the dark, followed by washing as described above. Membranes were air-dried on paper towels, and fluorescence signals were detected using an Amersham Typhoon scanner (Cytiva).

## Acknowledgements

We thank M. Hashimoto for her great technical assistance. We are grateful to the members of the Tamura laboratory for helpful discussion.

## Author Contributions

Conceptualization, Y.T.; Methodology, A.M., S.F., and Y.T.; Investigation, A.M. and Y.T.; Writing – Original Draft, Y.T.; Writing – Review & Editing, A.M., S.F., and Y.T.; Funding Acquisition, Y.T.; Supervision, Y.T.

