## Supplementary material for "A nucleus-vacuole junction in fission yeast enriches the HMG-CoA reductase Hmg1 and INSIG protein Ins1": Fig. S1

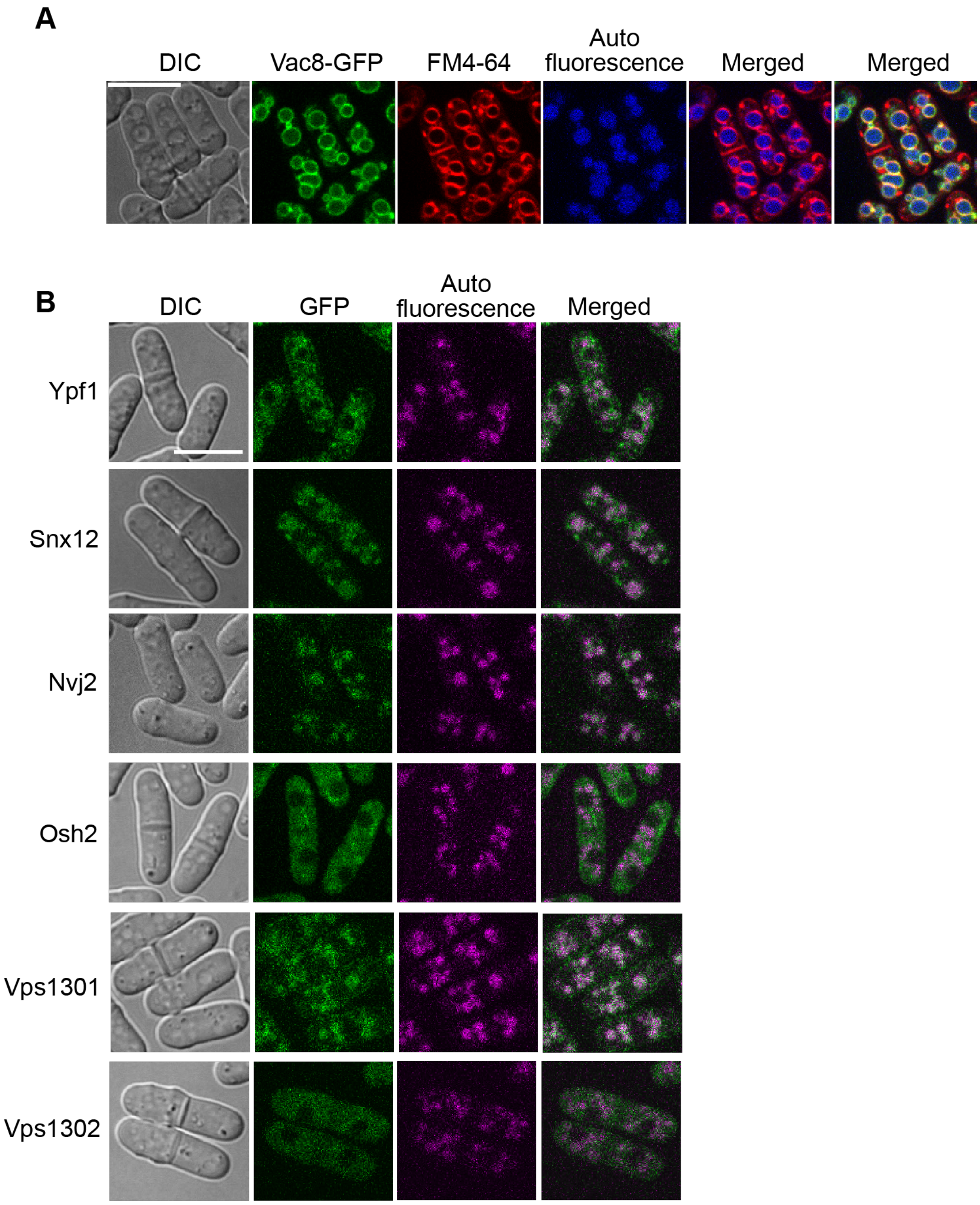


**Fig. S1. Hmg1 and Ins1 are enriched at the NVJ in fission yeast.**
(A) Cells expressing Vac8-GFP were stained with FM4-64 and observed by confocal microscopy. Scale bar, 10 µm. (B) Fission yeast orthologs of NVJ-localized proteins in budding yeast were tagged with GFP and observed by fluorescence microscopy. Vacuoles were visualized by autofluorescence. Scale bar, 10 µm.
