## Supplementary material for "A nucleus-vacuole junction in fission yeast enriches the HMG-CoA reductase Hmg1 and INSIG protein Ins1": Table S1

Table S1. Primers used in this study.

| Identifier 1 | Identifier 2 | Sequence |
|---|---|---|
| 5698 | spYpf1-tag-F | GTGACGAATTGAAAACACTCTTCTCCTTTCGTAGTGAAACAGAAGATGAAACTGACGAACAAGATAAATGTAAAAGTACTcggatccccgggttaattaa |
| 5699 | spYpf1-tag-R | TATGGCGAATAACGGATGAGCGTGAAGTAAGGAATGGGTTTTAACAAACGAATGTTGAAATTCTAAATCAAGGACCAATTgaattcgagctcgtttaaac |
| 5700 | spYpf1-tag-check-F | CCTCCCAGCCTAAGAAGCACTCG |
| 5701 | spYpf1-check-R | CGGATGAGCGTGAAGTAAGGAATGGG |
| 5702 | spNvj2-tag-F | CCCACAAACCTTTACCTAGGCCTCCCGTACAGGTTGAAACACGAGAGCCGGTTCGACCTGTACCGCCTATACCAAAACTTcggatccccgggttaattaa |
| 5703 | spNvj2-tag-R | TATAAGAAATGTGAGAACTTCTTATCAAAAGGATTGTGTCAAGGGAGGCCAAAGATACACTTTTGTATAAAGCCTTAGTGgaattcgagctcgtttaaac |
| 5704 | spNvj2-tag-check-F | GTAGCTCCAATAGCCCATCTG |
| 5705 | spNvj2-tag-check-R | GGATTGTGTCAAGGGAGGCC |
| 5706 | spIns1-tag-F | AAATTCAGTTTCGTTTATGGATTCCTATGATACTTTTCTCCGCTTCTACGATTGTTGGTAACGCTGGTCGTCTACTATTTcggatccccgggttaattaa |
| 5707 | spIns1-tag-R | TAACCCGGTTGTTTTGGATAATGAAGTCTAAGGCTAAAAAGAGAACGTAAAGCACAAACAACCAACTCCTAGAGAATAAAgaattcgagctcgtttaaac |
| 5708 | spIns1-tag-check-F | GCACTCGTTGCGAGTTCTGCTAGC |
| 5709 | spIns1-tag-check-R | CGCACACAACGATGAATGAGACG |
| 5710 | spHmg1-tag-F | ATACTCCTGCTATGGACTCTTCTGCCAAGAAACCAGCGACTGATGCTCTAAAATCCGTTAACTCTCGAGTACCGGGACGTcggatccccgggttaattaa |
| 5711 | spHmg1-tag-R | TAACTTTTTAGGGGAGATCAAAATTAAAAAAATTTGTCATCATCGTGTATGTGACAATATTATAAAACAAGCTTGCTCCCgaattcgagctcgtttaaac |
| 5712 | spHmg1-tag-check-F | CGTTGCAGCCGCCGTAATGGCT |
| 5713 | spHmg1-tag-check-R | CATTGCCTTACGTCTTCTTACC |
| 5941 | spINS1-del-F | ACGTAAAGATTGATTCTTCTCCATCCATTTAGTCAGTTTCACAAGGATCTCTACAGAAAGGTGCCGTAAGCTTGAATAAAgttgtaaaacgacggccagt |
| 5942 | spINS1-del-R | TAACCCGGTTGTTTTGGATAATGAAGTCTAAGGCTAAAAAGAGAACGTAAAGCACAAACAACCAACTCCTAGAGAATAAAcacaggaaacagctatgacc |
| 5943 | spINS1-check-F | CTCTACAGAAAGGTGCCGTAAG |
| 5944 | spINS1-check-R | CCCGGTTGTTTTGGATAATG |
| 5945 | spYPF1-del-F | TGAGGAAATATTTTCACGAGTATTTGAGGTTTGAAGGGGATAAGTTTTTGGATTTCCTTTGGTTCATCATTTTACAAAAGgttgtaaaacgacggccagt |
| 5946 | spYPF1-del-R | TATGGCGAATAACGGATGAGCGTGAAGTAAGGAATGGGTTTTAACAAACGAATGTTGAAATTCTAAATCAAGGACCAATTcacaggaaacagctatgacc |
| 5947 | spYPF1-check-F | CTGCTTGTTGCATCAGGTTACC |
| 5948 | spYPF1-check-R | GTATGGCGAATAACGGATGAGCG |
| 5949 | spVAC8-del-F | TTTGCTAACTGTCCTTGCAATTTGCTTTTACTGTCACTTAACTTCCTCCAAACACTAAACAATCATATAAACAAAGTCAGgttgtaaaacgacggccagt |
| 5950 | spVAC8-del-R | TTCCGAACTGGCAAACGGGGTTGAGACGGTAAAAATATAAAGAGATGAAGGGTAATATAATGTGCCAATAAAAATAAAAAcacaggaaacagctatgacc |
| 5951 | spVAC8-check-F | TGGATAACACAGCAACCCTG |
| 5952 | spVAC8-check-R | CTGGCAAACGGGGTTGAGACGG |
| 5953 | spVAC8-tag-F | GAGAGTATGAAGATGGTGAAGGCGATGTTATATTATTATCTGGCCGGGCATTGCATTTAATTGAACAGGACACAGATTCTcggatccccgggttaattaa |
| 5954 | spVAC8-tag-R | TTCCGAACTGGCAAACGGGGTTGAGACGGTAAAAATATAAAGAGATGAAGGGTAATATAATGTGCCAATAAAAATAAAAAgaattcgagctcgtttaaac |
| 5955 | spVAC8-tag-check-F | GGCAACTCTGCGGCTGCTCTGGG |
| 5977 | Snd302-tag-F | CCAATTCTAACAATGCTTCAGCTAAGTCTGATGGCCCTACAATCACTGAATTGAACGAAAACGAAACTGAAAAGTCTTCTcggatccccgggttaattaa |
| 5978 | Snd302-tag-R | TCAACCAGTGAAAGGACGAAATAATTAAACAAGCATGATGACTTTAGAAGTATCATTGAATGTCCAGCGTTCATAGATACgaattcgagctcgtttaaac |
| 5979 | Snd302-del-F | TCTAAGTTGTCCTCCACCATTACCTAAAAAAGAGGCAGTTGTAAGCTAAGTTATTCTTTTTTATTTCTAAAATTCTGAAAgttgtaaaacgacggccagt |
| 5980 | Snd302-del-R | TCAACCAGTGAAAGGACGAAATAATTAAACAAGCATGATGACTTTAGAAGTATCATTGAATGTCCAGCGTTCATAGATACcacaggaaacagctatgacc |
| 5981 | Snd302-check-F | AAGTTGTCCTCCACCATTACC |
| 5982 | Snd302-check-R | CTGGTTTGTGTAACACTTGAGC |
| 5983 | Snd302-tag-check-F | GGAGGAAACAAGCCTGCAGCTG |
| 5984 | NdeI-SpIns1-F | aattcatATGAGCAGAAAAGAGATTTACG |
| 6118 | pDUAL-FFH-spSND302-start-R | ATCCAGGCCTGTCGAAGAAGACTTTTCAGTTTCGTTTTCG |
| 6119 | pDUAL-seq-F | CTAGAGCAAAACACTAACCGCCA |
| 6120 | FFH-seq-R | GGTGATGTTTATCATCGTCGTC |
| 6156 | M13/pUC_Reverse | AGCGGATAACAATTTCACACAGG |
| 6157 | spLEU1_integration_R | CACAGCGACAACTCGGTCAT |
| 6236 | spSSP2-del-F | GGAAAAAAAGATTTTATCAATAAACTTTCAATATTGGATTTGCCAAGCGAATAGCTGATAgttgtaaaacgacggccagt |
| 6237 | spSSP2-del-R | CAACAATGGGTAATTAAGGGATGCTTGAAAACAGCGTGTTAAATGGGGAGTTACAATAAAcacaggaaacagctatgacc |
| 6238 | spSSP2-check-F | ACGCCGTGTGTTCCATTATA |
| 6239 | spSSP2-check-R | TGGCGGTAATTAATCCGGAC |
| 6240 | spNVJ2-del-F | TACTAAAAACATAAACACCAAGATCTGCTTTATTGAACTCTTCATGACTTTATTTTAATAgttgtaaaacgacggccagt |
| 6241 | spNVJ2-del-R | CTTATCAAAAGGATTGTGTCAAGGGAGGCCAAAGATACACTTTTGTATAAAGCCTTAGTGcacaggaaacagctatgacc |
| 6242 | spNVJ2-check-F | TTGAAACCATACGCTAGAAT |
| 6243 | spNVJ2-check-R | CTAGTGTTTCAGTTTGAACG |
| 6244 | spSND301-tag-F | CAAGTTTTGACTGATAAAAAAAGTGTGGATGAAGCTGAATTAACCAAACTCAAGCCCACTcggatccccgggttaattaa |

| | | |
|---|---|---|
| 6245 | spSND301-tag-R | AGGTGTGTTGTGGAAGAAGAGAAATTATTAAAAATTCCTGACAACTTAACAACACATTTTgaattcgagctcgtttaaac |
| 6246 | spSND301-del-F | TATTATCAACACCAAATGTATGAGAGGTAGTCTGAGTTCAACGTCTACACTTCGTACAAAgttgtaaaacgacggccagt |
| 6247 | spSND301-del-R | AGGTGTGTTGTGGAAGAAGAGAAATTATTAAAAATTCCTGACAACTTAACAACACATTTTcacaggaaacagctatgacc |
| 6248 | spSND301-check-F | GCCAAGTTGGATCAGATAGA |
| 6249 | spSND301-check-R | AGCTTTTAATGAAACGCGAT |
| 6250 | spTSC13-tag-F | TATCTGAAAGAGTTTCCCAATTATCCCCGTTCTCGTAAAATTATGATTCCCTTCTTTTTAcggatccccgggttaattaa |
| 6251 | spTSC13-tag-R | GATCAACATCTTTAAACTGATCGATCAAGAGTGAAAATCGATTGGTTCTAAATTTATAGAgaattcgagctcgtttaaac |
| 6252 | spTSC13-check-F | TCTAATCGAGGCTGTCTCTG |
| 6253 | spTSC13-check-R | GCTCAACAATTGTCTAGAGC |
| 6254 | spLTC1-tag-F | GAGCGAAGACTTAAAAAGCTAAAGGAAAGACTTAGAAAACTAGAGGCAAGTGGATATATTcggatccccgggttaattaa |
| 6255 | spLTC1-tag-R | AGCCCTGACGAAAAAAGCAGGAAAAATACATTGAAAATTCAGTCGCCGAATCAACCGATGgaattcgagctcgtttaaac |
| 6256 | spLTC1-del-F | ATTCATCGATTTAATACGTAGCTTCTTTTTGGTTAAGATCTGACAATTACCATACTTTAGgttgtaaaacgacggccagt |
| 6257 | spLTC1-del-R | AGCCCTGACGAAAAAAGCAGGAAAAATACATTGAAAATTCAGTCGCCGAATCAACCGATGcacaggaaacagctatgacc |
| 6258 | spLTC1-check-F | GAAAGAATCAGGTTACCACT |
| 6259 | spLTC1-check-R | ATTATTCGTGAAGCAACATG |
| 6260 | spSNX12-tag-F | CTTCTTGACGAGATTTTGCTGGCTTTAAAGAAACACGCTAGATCAACTAATAAGGCAACTcggatccccgggttaattaa |
| 6261 | spSNX12-tag-R | GAAAAGCATGAACATGATTACTCTTGCTATGATTTAACAAGAGAGGTAAAAAAGTGACACgaattcgagctcgtttaaac |
| 6262 | spSNX12-del-F | ATTTTTCCAACTAGAAAATCCATAGATAGAAGGACTTGGCTCTTACCATTATAGGACCAAgttgtaaaacgacggccagt |
| 6263 | spSNX12-del-R | GAAAAGCATGAACATGATTACTCTTGCTATGATTTAACAAGAGAGGTAAAAAAGTGACACcacaggaaacagctatgacc |
| 6264 | spSNX12-check-F | CTTTGCTTACCATTTTCCGG |
| 6265 | spSNX12-check-R | CTGGCCGTTAGAAGATAAAC |
| 6266 | spSNX12-tag-check-F | TCATGATCACTCCAAAGACC |
| 6267 | spVps1301-tag-F | ATTCGAGACGAGCTTGCCGCTTATAAACACAAAGTAAATTACGAGCTAGAGGTTGCTCTTcggatccccgggttaattaa |
| 6268 | spVps1301-tag-R | ACAAAACATTAATATATACAAATTCCTTCATAAGAGTTCAATAACAAATAAACAAAAAAAgaattcgagctcgtttaaac |
| 6269 | spVps1301-tag-check-F | GCTAAGATTGTCAAGACTGA |
| 6270 | spVps1301-check-R | AAGGCAAGCCATCGTTATCT |
| 6479 | spSnd302-147aa-tag-F | TTGATTTCCTTGTTTACTAACAACTTGGTTTCCATTTACATCTTTAATAAGGCTGCTGAAcggatccccgggttaattaa |
| 6480 | spSnd302-159aa-tag-F | TACATCTTTAATAAGGCTGCTGAAGGTAGCTTATCTCGTCCATTTGCTCCCCCTGCTGGTcggatccccgggttaattaa |
| 6481 | spSnd302-182aa-tag-F | GGAGGAAACAAGCCTGCAGCTGCTGTTACTGGCACTTCGTCCAATTCTAACAATGCTTCAcggatccccgggttaattaa |
| 6493 | Tadh-SLiCE-ADEL-R | cgcttatttagaagtttaAAGTTCATCGGCCTCATCATCGAAGTACTTGTACAGCTCGTCCATGC |
| 6494 | mCherry_spBip1_35aa_R | tcctcgcccttgctcaccatTCCATATGATTCTGTAGAGTTATC |
| 6495 | spBip1_35aa_mCherry_F | GATAACTCTACAGAATCATATGGAatggtgagcaagggcgagga |
| 6496 | Pcam1-SLiCE-spBip1-F | tttagagttactgatATGAAGAAGTTCCAGCTATT |
| 6497 | Pef1a-SLiCE-spSnd302-F | tttcatcgaactcacATGAACGCTCAGGTATTAAATCTTGTCGCCGCATTGGG |
| 6578 | spOSH2-tag-F | GCTGGTGAAGCTCATTTGGCCGGAAAGGAATTTGAATGGCCTAATGTTGATGATATTTTTcggatccccgggttaattaa |
| 6579 | spOSH2-tag-R | AGTTGGATTGCCATTCCTTGCCTTTTTGCAAAACAAAATGATTAATAATTTCGTGAGATCgaattcgagctcgtttaaac |
| 6582 | spOSH2-check-F | AACCATAAGTGATGAACCGA |
| 6583 | spOSH2-check-R | CAGTGTTGTTGGTCATAGTG |
| 6711 | spVps1302-tag-F | GCTGCATTTACTGCTTATTCAGAACATTATAGATTGCGATTGAAGCAGATATCCCCACTTcggatccccgggttaattaa |
| 6712 | spVps1302-tag-R | ATAGAAGAAAATATACAATAGTTTAAAAAGTAAACAAAAATATAAAAAAAAATTGATAATgaattcgagctcgtttaaac |
| 6713 | spVps1302-del-F | TCTGTAAAGCAAAGTTCAAAATATTTACAGTTTGCTGTTTCTCACCGTATTCCCTTCAATgttgtaaaacgacggccagt |
| 6714 | spVps1302-del-R | ATAGAAGAAAATATACAATAGTTTAAAAAGTAAACAAAAATATAAAAAAAAATTGATAATcacaggaaacagctatgacc |
| 6715 | spVps1302-check-F | TTACTTATGGTGTAGGCGTT |
| 6716 | spVps1302-check-R | CAGTCAAAGAGGGATTCACA |
| 6717 | spVps1302-tag-check-F | TTTTGAAGGGATTTGGCAAG |
| 6732 | spTsc13-del-F | CCACGAGCTTCTTAAGCCCAACAACAACAGCACACAGCTTGTTAAAGGTAGGTTTTCAAAgttgtaaaacgacggccagt |
| 6733 | spTsc13-del-R | GATCAACATCTTTAAACTGATCGATCAAGAGTGAAAATCGATTGGTTCTAAATTTATAGAcacaggaaacagctatgacc |
| 6768 | Pef1a-SLiCE-ScSND3-F | tttcatcgaactcacATGAATCCTCAAGTCAGTAA |
| 6770 | Tadh1-SLiCE-ScSND3-R | cgcttatttagaagttcaTTCAGCCTTAACACCAG |
| 6771 | Pef1a-SLiCE-SpSND301-F | tttcatcgaactcacATGGTTTCAAGATGGGAGAA |
| 6772 | Tadh1-SLiCE-SpSND301-R | cgcttatttagaagtttaAGTGGGCTTGAGTTTGG |
| 6773 | spElo1-del-F | TTTTCGTTACTTGCTAGTCTCCGCCATTTGTACCCTGCAAATCTCAAATACGTTTTCATTgttgtaaaacgacggccagt |
| 6774 | spElo1-del-R | CAAGTTATAAGTAAGAGCAAAATCGGGTAATTAAAAACCATAAGAGGTACCACGAAAACAcacaggaaacagctatgacc |
| 6775 | spElo1-check-F | TTTGTGCAAATATCACAACG |
| 6776 | spElo1-check-R | CGTAAGTCACTCCAACTATT |
