## Supplementary material for "A nucleus-vacuole junction in fission yeast enriches the HMG-CoA reductase Hmg1 and INSIG protein Ins1": Table S2

Table S2. Fission yeast strains used in this study

| Identifier | name | Genotype |
| --- | --- | --- |
| 1C6 | WT | *h90 ade6-216 leu1-32 lys1-131 ura4-D18* |
| 1A3 | *ypf1-GFP* | *h90 ade6-216 leu1-32 lys1-131 ura4-D18 ypf1-GFP::KanMX* |
| 1A6 | *nvj2-GFP* | *h90 ade6-216 leu1-32 lys1-131 ura4-D18 nvj2-GFP::KanMX* |
| 1A9 | *ins1-GFP* | *h90 ade6-216 leu1-32 lys1-131 ura4-D18 iins1-GFP::KanMX* |
| 1B2 | *hmg1-GFP* | *h90 ade6-216 leu1-32 lys1-131 ura4-D18 hmg1-GFP::KanMX* |
| 1D3 | *ins1-GFP hmg1-mCherry* | *h90 ade6-216 leu1-32 lys1-131 ura4-D18 iins1-GFP::KanMX hmg1-mCherry::Hyg* |
| 1D4 | *vac8-GFP* | *h90 ade6-216 leu1-32 lys1-131 ura4-D18 vac8-GFP::KanMX* |
| 1E2 | *snd302-GFP* | *h90 ade6-216 leu1-32 lys1-131 ura4-D18 snd302-GFP::KanMX* |
| 1E6 | *snd302-3×FLAG* | *h90 ade6-216 leu1-32 lys1-131 ura4-D18 snd302-3×FLAG::KanMX* |
| 1E8 | *ins1-GFP snd302Δ* | *h90 ade6-216 leu1-32 lys1-131 ura4-D18 iins1-GFP::KanMX snd302Δ::natNT2* |
| 1G2 | *hmg1-GFP ypf1Δ* | *h90 ade6-216 leu1-32 lys1-131 ura4-D18 hmg1-GFP::KanMX ypf1Δ::natNT2* |
| 1G4 | *hmg1-GFP vac8Δ* | *h90 ade6-216 leu1-32 lys1-131 ura4-D18 hmg1-GFP::KanMX vac8Δ::natNT2* |
| 1G5 | *hmg1-GFP snd302Δ* | *h90 ade6-216 leu1-32 lys1-131 ura4-D18 hmg1-GFP::KanMX snd302Δ::natNT2* |
| 1G7 | *hmg1-GFP: ins1Δ* | *h90 ade6-216 leu1-32 lys1-131 ura4-D18 hmg1-GFP::KanMX ins1Δ::natNT2* |
| 1I3 | *hmg1-GFP ltc1Δ* | *h90 ade6-216 leu1-32 lys1-131 ura4-D18 hmg1-GFP::KanMX ltc1Δ::natNT2* |
| 1I5 | *hmg1-GFP snd301Δ* | *h90 ade6-216 leu1-32 lys1-131 ura4-D18 hmg1-GFP::KanMX snd301Δ ::natNT2* |
| 1I7 | *hmg1-GFP snx12Δ* | *h90 ade6-216 leu1-32 lys1-131 ura4-D18 hmg1-GFP::KanMX snx12Δ::natNT2* |
| 1J10 | *hmg1-GFP ssp2Δ* | *h90 ade6-216 leu1-32 lys1-131 ura4-D18 hmg1-GFP::KanMX ssp2Δ::natNT2* |
| 2B4 | *hmg1-GFP snd302(FL)-FLAG* | *h90 ade6-216 leu1-32 lys1-131 ura4-D18 hmg1-GFP::KanMX snd302 (FL)-6×Gly-FLAG::hphMX* |
| 2B5 | *hmg1-GFP snd302(1-147aa)-FLAG* | *h90 ade6-216 leu1-32 lys1-131 ura4-D18 hmg1-GFP::KanMX snd302 (1-147aa)-6×Gly-FLAG::hphMX* |
| 2B7 | *hmg1-GFP snd302(1-159aa)-FLAG* | *h90 ade6-216 leu1-32 lys1-131 ura4-D18 hmg1-GFP::KanMX snd302 (1-159aa)-6×Gly-FLAG::hphMX* |
| 2B9 | *hmg1-GFP:: snd302(1-182aa)-FLAG* | *h90 ade6-216 leu1-32 lys1-131 ura4-D18 hmg1-GFP::KanMX snd302 (1-182aa)-6×Gly-FLAG::hphMX* |
| 2C1 | *snx12-mNeonGreen-3×FLAG* | *h90 ade6-216 leu1-32 lys1-131 ura4-D18 snx12-mNeonGreen-3×FLAG::hphMX* |
| 2C4 | *vps1301-mNeonGreen-3×FLAG* | *h90 ade6-216 leu1-32 lys1-131 ura4-D18 vps1301-mNeonGreen-3×FLAG::hphMX* |
| 2C6 | *ltc1-mNeonGreen-3×FLAG* | *h90 ade6-216 leu1-32 lys1-131 ura4-D18 ltc1-mNeonGreen-3×FLAG::hphMX* |
| 2C7 | *tsc13-mNeonGreen-3×FLAG* | *h90 ade6-216 leu1-32 lys1-131 ura4-D18 tsc13-mNeonGreen-3×FLAG::hphMX* |
| 2C10 | *osh2-mNeonGreen-3×FLAG* | *h90 ade6-216 leu1-32 lys1-131 ura4-D18 osh2-mNeonGreen-3×FLAG::hphMX* |
| 2D1 | *snd301-mNeonGreen-3×FLAG* | *h90 ade6-216 leu1-32 lys1-131 ura4-D18 snd301-mNeonGreen-3×FLAG::hphMX* |
| 2D4 | *snd301-3×FLAG* | *h90 ade6-216 leu1-32 lys1-131 ura4-D18 snd301-3×FLAG::KanMX* |
| 2E1 | *hmg1-GFP snd302Δ Pef1a-snd302-2×FLAG-6×His-Tadh1* | *h90 ade6-216 leu1-32 lys1-131 ura4-D18 hmg1-GFP::KanMX snd302Δ::natNT2 Pef1a-snd302-3×FLAG-6×His-Tadh1::leu1* |
| 2G3 | *vps1302-mNeonGreen-3×FLAG::hphMX* | *h90 ade6-216 leu1-32 lys1-131 ura4-D18 vps1302-mNeonGreen-3×FLAG::hphMX* |
| 2H8 | *hmg1-GFP nvj2Δ* | *h90 ade6-216 leu1-32 lys1-131 ura4-D18 hmg1-GFP::KanMX nvj2Δ::natNT2* |
| 3A1 | *hmg1-GFP Pcam1-BipN-mCherry-ADEL-Tadh1* | *h90 ade6-216 leu1-32 lys1-131 ura4-D18 hmg1-GFP::KanMX Pcam1-BiipN-mCherry-ADEL-Tadh1::leu1* |
| 3B5 | *hmg1-GFP snd302Δ::natNT2 Pef1a-scsnd3-Tadh1* | *h90 ade6-216 leu1-32 lys1-131 ura4-D18 hmg1-GFP::KanMX snd302Δ::natNT2 Pef1a-scsnd3-Tadh1::leu1* |
| 3C3 | *hmg1-GFP snd302Δ Pcam1-BipN-mCherry-ADEL-Tadh1* | *h90 ade6-216 leu1-32 lys1-131 ura4-D18 hmg1-GFP::KanMX snd302Δ::natNT2 Pcam1-BiipN-mCherry-ADEL-Tadh1::leu1* |
| 3C4 | *hmg1-GFP snd302Δ Pef1a-snd301-Tadh1* | *h90 ade6-216 leu1-32 lys1-131 ura4-D18 hmg1-GFP::KanMX snd302Δ::natNT2 Pef1a-snd301-Tadh1::leu1* |
| 3D2 | *hmg1-GFP vps1302Δ* | *h90 ade6-216 leu1-32 lys1-131 ura4-D18 hmg1-GFP::KanMX vps1302Δ::natNT2* |
| 4A4 | *hmg1-GFP vps1301Δ* | *h90 ade6-216 leu1-32 lys1-131 ura4-D18 hmg1-GFP::KanMX vps1301Δ::natNT2* |
| 4A7 | *hmg1-GFP elo1Δ* | *h90 ade6-216 leu1-32 lys1-131 ura4-D18 hmg1-GFP::KanMX elo1Δ::natNT2* |
| 4C9 | *hmg1-GFP vac8-mCherry* | *h90 ade6-216 leu1-32 lys1-131 ura4-D18 hmg1-GFP::KanMX vac8-mCherry::Hyg* |
